# The collapse of genetic incompatibilities in hybridizing populations

**DOI:** 10.1101/2021.01.08.425971

**Authors:** Tianzhu Xiong, James Mallet

## Abstract

Diverging species are often genetically incompatible upon hybridization. Such incompatibilities are considered important in keeping the integrity of species from the disruption of hybrids. However, recent empirical work has shown that not all incompatibilities are gene-flow-proof, and they can collapse due to continuing hybridization. Counterintuitively, many studies found that incompatible alleles are already segregating within species, whereas they should go extinct quickly in a randomly mating population. Due to the complexity of multilocus epistasis, few general principles explain behaviors of incompatibilities under gene flow both within and between species. In the current work, we argue that the redundancy of genetic mechanisms can robustly determine the dynamics of intrinsic incompatibilities under gene flow. While higher genetic redundancy decreases the stability of incompatibilities during hybridization, it also increases the tolerance of incompatibility polymorphism within each species. We treat two general classes of incompatibilities. In the redundant class, similar to the classical Dobzhansky-Muller system, the collapse is continuous and eventually approaches quasi-neutral polymorphism between broadly-sympatric species, often as a result of isolation-by-distance. In the non-redundant class, analogous to the shifting-balance process, incompatibilities collapse abruptly with spatial traveling waves. We obtained simulated and analytical results for several incompatibility models to demonstrate the differences between the two classes. As both redundant and non-redundant genetic mechanisms of incompatibilities are common, the proposed conceptual framework may help understand the abundance of incompatibilities in natural populations.

## 1. Introduction

Speciation, the process by which multiple lineages arise from a common ancestor and persist through time, is often accompanied by increased incompatibility between genomes (Dagilis et al., 2019; Matute et al., 2010; Orr and Turelli, 2001). Although the evolution of such incompatibilities might not be the direct cause of initial divergence, their presence is considered important for maintaining species integrity when different lineages meet in sympatry (Coughlan and Matute, 2020). The conceptual picture of genetic incompatibilities has been largely explored using the classical two-locus Dobzhansky-Muller incompatibility model (Dobzhansky, 1937; Muller, 1942), where derived alleles arise on different loci in each species, and these alleles interact negatively to produce hybrids with lower fitness. We hereafter use the term “classical Dobzhansky-Muller systems” only as the synonym of the original version of such incompatibilities, although it has been often used broadly as the synonym of any epistatic incompatibilities.

Recent experimental characterization of genetic incompatibilities has revealed considerably more complex genetic architectures as well as population genetic patterns. While incompatibilities in hybrids are generally epistatic, multiple interacting loci often explain dysfunction in hybrids (complex epistasis) (Courret et al., 2019; Meiklejohn et al., 2018; Rosser et al., 2021). Incompatibility is also variable within species (Atlan et al., 2003; Corbett-Detig et al., 2013; Cutter, 2012; Larson et al., 2018; Zuellig and Sweigart, 2018b). In some cases, balancing selection has been invoked to explain widespread polymorphism (Seidel et al., 2008), but it is unclear if most incompatibility variation has reached long-term equilibria across space and time. Incompatibility genes could also flow among young lineages, as has been confirmed or suspected in a number of hybridizing systems (Larson et al., 2018; Presgraves and Meiklejohn, 2021; Rosser et al., 2019). Since hybridization is a rather frequent phenomenon among recently diverged and incipient species (Mallet, 2005), understanding incompatibilities subject to gene flow is crucial for a more complete picture of incompatibility variation in nature.

To date, a considerable number of theories have been proposed to deal with the accumulation of incompatibility alleles between a pair of isolated, panmictic populations (Dagilis et al., 2019; Orr and Turelli, 2001; Schiffman and Ralph, 2021), where polymorphism is viewed as a transient phenomenon of little importance. In these “accumulation” theories, hybridization at most resembles laboratory crosses with little impact on parental species. On the other hand, theoretical analysis of incompatibility dynamics under gene flow has hitherto been limited to a single population receiving migrants from a large, static species, often assuming no genetic drift (Bank et al., 2012; Blanckaert et al., 2020; Blanckaert and Hermisson, 2018). The complexity of incompatibility systems in natural populations is likely much higher, especially if non-equilibrium patterns from hybridization and population structures are considered. Of particular interest are patterns generated by intrinsic properties of incompatibilities (i.e., no ecological selection), because intrinsic properties are more likely to be shared among different taxa with a similar genetic basis of incompatibilities, while ecological selection can always be added as an extra layer when necessary. If such intrinsic patterns exist and change only slowly through time, they might account for a nontrivial component of observed data.

Nonetheless, many incompatibilities are polygenic and epistatic, which obstructs the formulation of simple, robust principles that are independent of model details. Therefore, we first present the intuition behind a general model underlying our technical analysis, that patterns of incompatibilities under hybridization are largely determined by the level of genetic redundancy. Using this guideline, we show qualitative differences of incompatibility collapse by analyzing and simulating several incompatibility systems under different population structures.

## 2. Models

### 2.1 Intuition: the duality between genetic redundancy and the stability of incompatibilities to hybridization

Redundant components such as duplicated genes or pathways greatly facilitate the evolution of genetic incompatibilities, as they produce extra degrees of freedom even for species under stabilizing selection (Haag, 2007; Lynch and Force, 2000). In theory, the classical Dobzhansky-Muller system could represent the simplest form of redundant incompatibilities, because substituting a single derived allele into the ancestral background has no fitness cost, showing that the fitness of ancestral genotype is robust against small-scale disruptions. However, a greater redundancy also de-stabilizes incompatibilities during hybridization, which is first shown in the classical Dobzhansky-Muller system that continuing hybridization could lead to its total collapse (Barton and Bengtsson, 1986). Empirical examples of redundant incompatibilities are not rare. For instance, in *Mimulus*, duplicated genes with reciprocal pseudogenization in each species create pseudogene-pseudogene incompatibilities (Zuellig and Sweigart, 2018a), which is typically unstable to hybridization (a particular case of the Dobzhanskly-Muller model) (Bank et al., 2012). In *Arabidopsis thaliana*, the existence of additional gene copies as rescuers of a known pseudogene-pseudogene incompatibility further increases redundancy and two pseudogenes could co-exist at arbitrary frequencies in a population (Jiao et al., 2021). Among other populations of *A. thaliana*, first-generation hybrid necrosis was mapped to a large array of dominantly interacting immune-response genes, which also arise via multiple cycles of gene duplication and gene loss (Chae et al., 2014). In chromosomal incompatibilities, karyotypic lineages of *Sorex araneus* with different Robertsonian fusions collapse into fully acrocentric and globally compatible chromosomes in the hybrid zone, resembling a karyotypic version of the Dobzhansky-Muller system (Hatfield et al., 1992). In regulatory networks, genetic redundancy also frequently appears in both *trans* and *cis* regulatory elements that could be important for the development of incompatibilities. For instance, distinct transcription factors could respond to similar signals, and control a similar set of target genes (AkhavanAghdam et al., 2016; Hu et al., 2007), and some enhancers are also functionally redundant (Barolo, 2012). In principle, if a biological function is redundantly controlled by several independent genetic modules, it may be robust to disruptions of a subset of them. Thus, even if all modules are divergent between species and might be disrupted in hybrids, natural selection will select for genotypes of at least one module and maintain fitness during hybridization, while all the other modules are freely exchanged between species, which de-stabilizes incompatibilities.

Conversely, incompatibilities lacking any redundancy are typically more resistant to hybridization. A classic example is the co-evolution between *cis* and *trans* acting elements, where mis-regulation almost always occurs in recombined genotypes (Barriere et al., 2012; Mack and Nachman, 2017; Porter and Johnson, 2002; Prager and Wilson, 1975). More generally, incompatibilities involving some form of reciprocal sign epistasis (Poelwijk et al., 2011), with examples such as gene-dosage imbalance (Josefsson et al., 2006), cyto-nuclear interactions (Chou and Leu, 2010), or selfish genetic elements with their suppressors (Case et al., 2016; Phadnis and Orr, 2009), are also typically non-redundant and could be bi-stable. Interestingly, some incompatibilities might be partially redundant. For instance, polygenic threshold incompatibilities are frequently observed in *Drosophila* (Liénard et al., 2016; Morán and Fontdevila, 2014), where strong hybrid dysfunctions occur only when the amount of introgressed DNA exceeds a certain fraction, so that these systems might tolerate a mild level of free-segregation involving incompatibility genes (Morimoto et al., 2020).

We emphasize that duplicated components in genomes change over time (Cotton and Page, 2005; Moore and Purugganan, 2005), as does genetic redundancy. Thus, incompatibility factors evolved by redundant processes might lack any redundancy when hybridization occurs. Consequently, the proposed conceptual duality should be between the *current* level of genetic redundancy and the stability of genetic incompatibilities to hybridization. This duality forms the basis of our models below.

### 2.2 Two incompatibility classes

We start by considering two genetically incompatible species, 1 and 2. Let Ω be the space of all possible genotypes that can be created by recombination and union of all haplotypes which pre-exist in either species 1 or species 2. The set of all probability distributions on Ω is denoted by G, which corresponds to the space of genotype distributions. G is the primary state space of the general model. Elements in G are denoted by **g**. We use **g**_1_ and **g**_2_ as the genotype distributions in the two species, respectively. We assume that generations are overlapping, and microscopic changes are governed by an individual-based birth-death process (see Materials and Methods: Construction of dynamics from a birth-death process). If genetic drift is neglected, evolution in a single randomly mating population is driven by viability selection and reproduction, the joint action of which induces a continuous-time dynamical system on G (A glossary of symbols and variables can be found in Table 1).

**Table 1:**
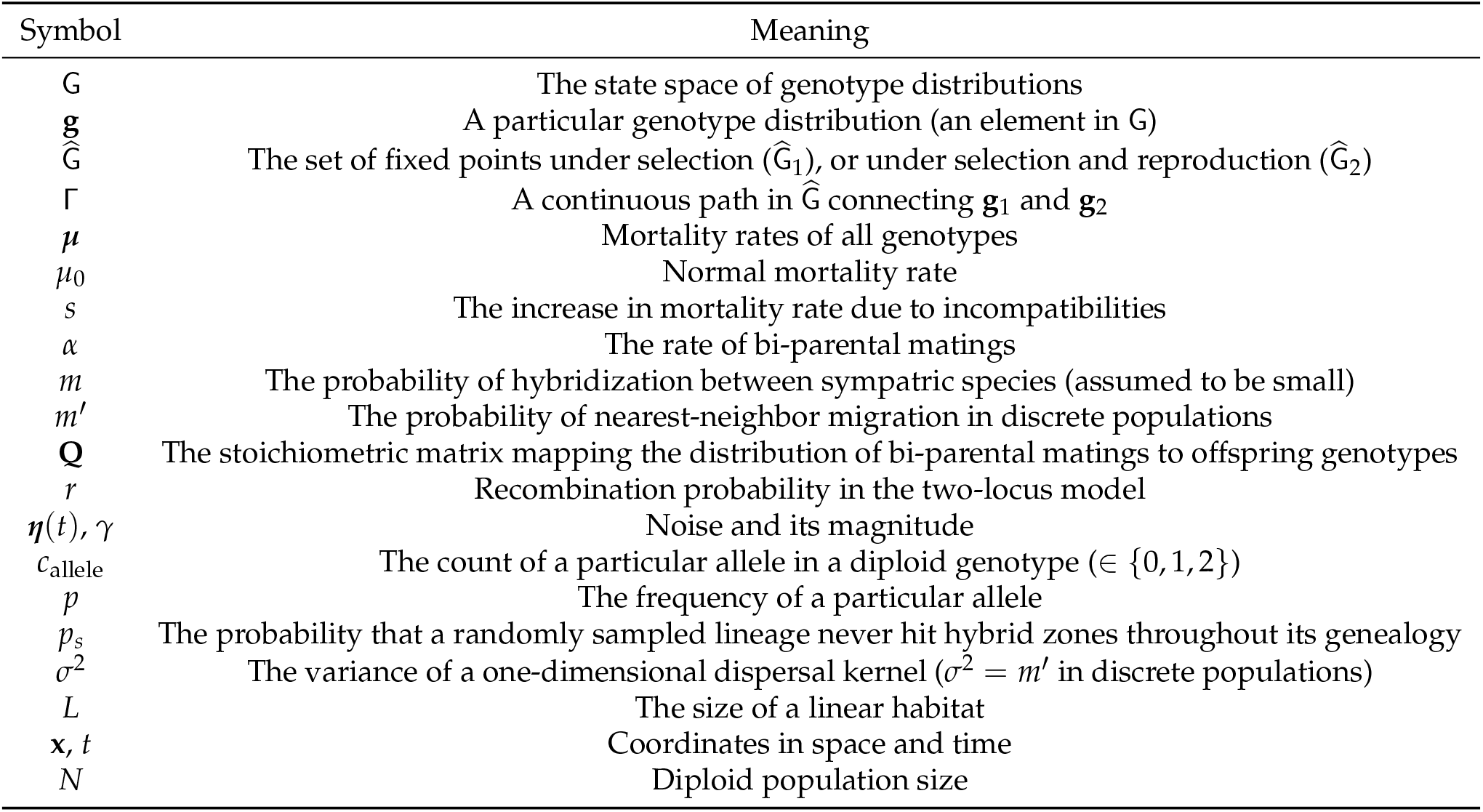
Glossary of major symbols

| Symbol | Meaning |
| --- | --- |
| $G$ | The state space of genotype distributions |
| $\mathbf{g}$ | A particular genotype distribution (an element in $G$) |
| $\hat{G}$ | The set of fixed points under selection ($\hat{G}_1$), or under selection and reproduction ($\hat{G}_2$) |
| $\Gamma$ | A continuous path in $\hat{G}$ connecting $\mathbf{g}_1$ and $\mathbf{g}_2$ |
| $\mu$ | Mortality rates of all genotypes |
| $\mu_0$ | Normal mortality rate |
| $s$ | The increase in mortality rate due to incompatibilities |
| $\alpha$ | The rate of bi-parental matings |
| $m$ | The probability of hybridization between sympatric species (assumed to be small) |
| $m'$ | The probability of nearest-neighbor migration in discrete populations |
| $\mathbf{Q}$ | The stoichiometric matrix mapping the distribution of bi-parental matings to offspring genotypes |
| $r$ | Recombination probability in the two-locus model |
| $\eta(t), \gamma$ | Noise and its magnitude |
| $c_{\text{allele}}$ | The count of a particular allele in a diploid genotype ($\in \{0, 1, 2\}$) |
| $p$ | The frequency of a particular allele |
| $p_s$ | The probability that a randomly sampled lineage never hit hybrid zones throughout its genealogy |
| $\sigma^2$ | The variance of a one-dimensional dispersal kernel ($\sigma^2 = m'$ in discrete populations) |
| $L$ | The size of a linear habitat |
| $\mathbf{x}, t$ | Coordinates in space and time |
| $N$ | Diploid population size |

An explicit construction of this dynamical system in a single isolated population can be written concisely as

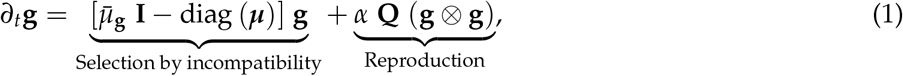

where *α* is the rate of offspring production via bi-parental mating, **Q** is the stoichiometric matrix relating the distribution of offspring genotypes to the distribution of bi-parental matings (given by the tensor product **g** ⊗ **g**), I is the identity matrix, vector *μ* contains the mortality rates of all genotypes, and its average value over a genotype distribution **g** is denoted as 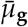. We assume the holey-landscape fitness model (Gavrilets, 1997) and ecological selection among fit genotypes is not considered as we are only interested in intrinsic properties of incompatibilities. Under the holey-landscape fitness model, mortality rates among genotypes are either normal or excessively high. To model the influence of hybridization from species 2 on species 1, split reproduction to include a small probability *m* of inter-specific hybridization, and we have the following equation governing the genotype distribution **g**_1_:

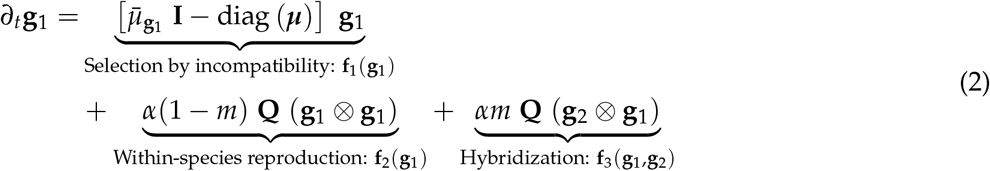

Note that the probability of interspecific gene flow under this formulation is *m*/2, because each first-generation hybrid only has half of its DNA inherited from its foreign parent. Consider the following sets of fixed points:

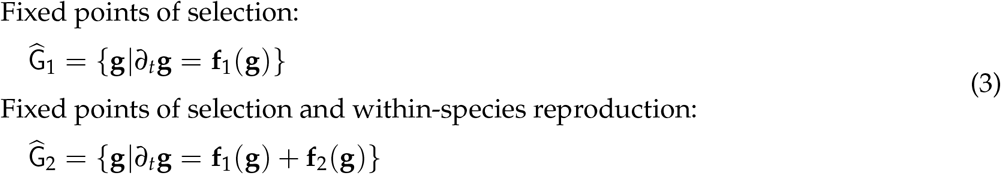

Since multiple genotypes are equally fit under the holey-landscape fitness model, both sets might become degenerate (i.e., not composed entirely of isolated points). This allows for the existence of continuous paths in both sets. If parental genotype distributions **g**_1_ and **g**_2_ are path-connected in either set, then it is possible to perturb **g**_1_ along such paths to **g**_2_ without leaving the equilibrium of some components of the dynamics in Equation (2). This is the dominant route of incompatibility collapse in our model.

Formally, denote a continuous path of fixed points connecting the parental genotype distributions by Γ_1_ in 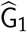, and by Γ_2_ in 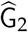 (Figure 1A). Showing the existence of Γ_1_ is trivial because the set {g|g = *λ***g**_1_ + (1 – *λ*)**g**_2_, (0 ≤ *λ* ≤ 1)} is invariant under the dynamics of **f**_1_. The existence of Γ_2_ is not always guaranteed, but its presence could be used as a *definition* of redundant incompatibilities, and so we get two different regimes of dynamics depending on whether genetic redundancy exists. Note that genetic redundancy is largely a qualitative notion, but it generally implies that maintaining high fitness does not require all components of the system to work (Nowak et al., 1997), which adds some free dimensions to the dynamics that are unconstrained by fitness. The existence of Γ2 is the consequence of such free dimensions^1^. Nonetheless, when Γ_2_ does not exist, dynamics retreat to the next-available path Γ_1_. This leads to the following two classes of incompatibilities.

**Figure 1:**
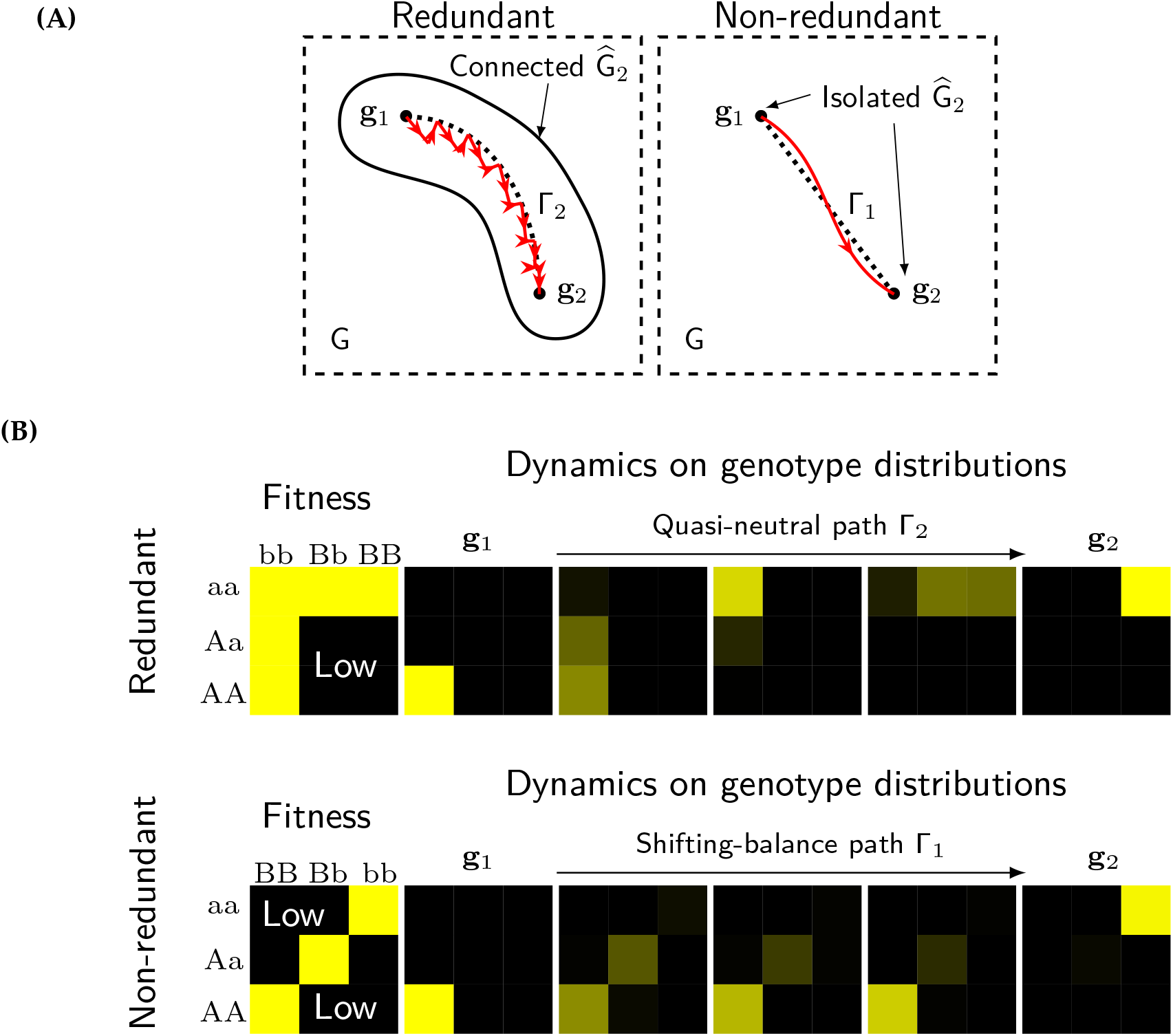
Schematic differences between the collapse dynamics of redundant and non-redundant incompatibilities. **(A)** Representation of dynamics in the general model. Red lines depict the trajectories of genotype distributions in species 1 subject to unidirectional gene flow from species 2. For redundant incompatibilities, their collapse trajectories closely follow a quasi-neutral path Γ_2_ in the connected set of fixed points 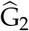. For non-redundant incompatibilities, their collapse is a large jump along Γ_1_, because fixed points under selection and reproduction are isolated. **(B)** Explicit examples of dynamics in two-locus models (see Section “Specific models”. Non-redundant: Model I; Redundant: Model II). The genotype distribution in species 1 is subject to unidirectional gene flow from species 2.

#### Non-redundant incompatibilities

If a quasi-neutral path Γ_2_ does not exist, under the holey-landscape fitness model, we can asymptotically eliminate only the time-scale of selection. Such systems are governed by (see Materials and Methods: Separation of time-scales):

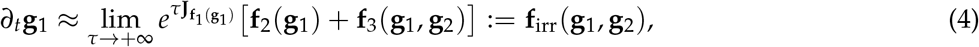

where **J** stands for the Jacobian matrix at equilibrium points of the dynamical forces specified by its subscript. Such incompatibilities are resolved along path Γ_1_ by an abrupt jump (i.e., shifting-balance. A two-locus example is given in Figure 1B).

#### Redundant incompatibilities

If a quasi-neutral path Γ_2_ exists, under suitable conditions we can asymptotically eliminate time-scales of selection and within-species reproduction, and the system evolves under the time-scale of hybridization:

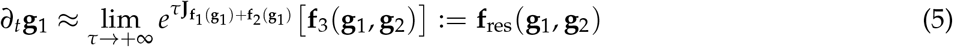

Such incompatibilities are often resolved along path Γ_2_ by incremental perturbations from hybridization (see Figure 1B for a two-locus example).

#### Noise

In a finite population, Equations (4) and (5) are driven by an additional noise term ***η***(*t*) encapsulating genetic drift and other stochastic effects in the birth-death process:

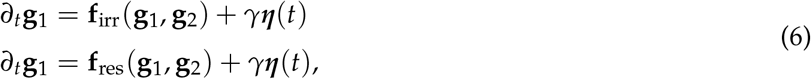

where *γ* is a scaling factor determining the magnitude of noise.

### 2.3 Specific models

Inspired by realistic molecular mechanisms, we here define four incompatibility models with different levels of redundancy (Table 2), on which all subsequent results will be based. Simpler models (I & II) are mainly used for the analytical treatment of collapse dynamics, and they have been constructed in existing literature (Lindtke and Buerkle, 2015). Complex models (III & IV) conceived for highly redundant incompatibilities are used for simulations in SLiM-3.6 (Haller and Messer, 2019) to demonstrate the robustness of certain quasi-neutral behaviors in spatially-extended populations. At most four bi-allelic loci in diploid species will be considered, and we track alleles that are represented by capitalized letters (A, B, C, D). Under the birth-death formulation, these models have the following form of mortality rates:

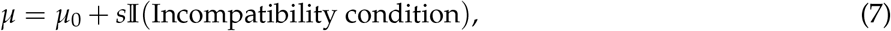

where 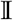 is the indicator function, *s* is the increase in mortality rates by incompatibility. Further, let *c*_allele_ be the count of the designated allele in a diploid genotype.

**Table 2:**
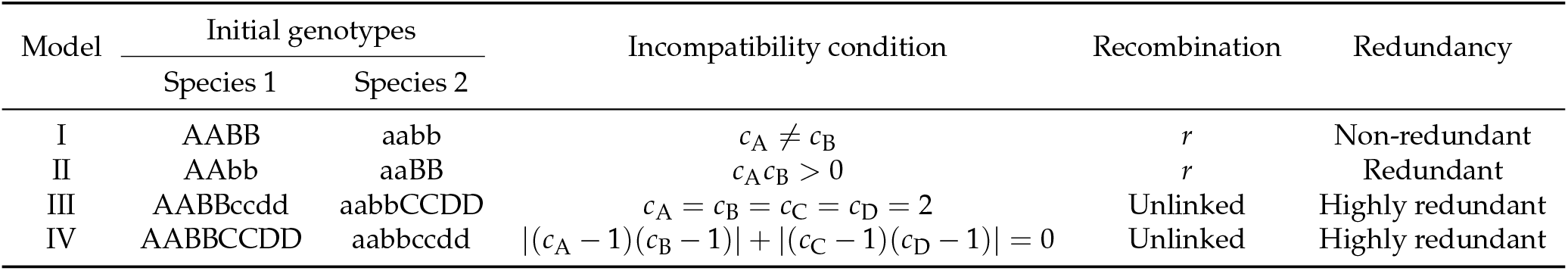
Specific models of incompatibilities.

#### Model I: two-locus bi-stable systems (non-redundant)

Bi-stable systems with two loci could represent the co-evolution between regulatory elements in a pathway under stabilizing selection. In the model, we assume fitness reduction is due to additive dosage-imbalance between two loci, which leads to incompatibility condition:

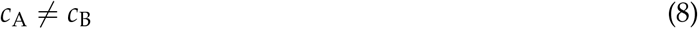

This bi-stable system broadly represents behaviors of non-redundant incompatibilities that are intrinsically symmetric, because mortality rates are invariant when exchanging labels “A” and “B”.

#### Model II: two-locus Dobzhansky-Muller systems (redundant)

The classical Dobzhansky-Muller system involves two derived alleles (A and B) on two loci, each behaving neutrally on its own. They interact negatively when combined in hybrids. For simplicity, negative epistasis is assumed to be fully dominant, so that incompatibility condition becomes

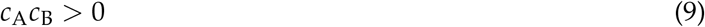

#### Model III: redundant coupling of uni-stable systems

This redundant system involves duplicating the same uni-stable locus. In the main text, we consider four pseudogenized alleles (A, B, C, D) of the same gene with four functional homologs (a, b, c, d). Each locus is uni-stable on its own because its functional allele will always replace the pseudogenized one. Mortality rate is normal whenever a functional copy is present in an individual, and incompatibility condition is

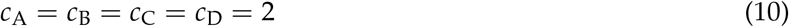

This model is also an extension to the two-locus Dobzhansky-Muller system with recessive interactions.

#### Model IV: redundant coupling of multi-stable systems

This system considers a type of genetic redundancy in pathway duplication, where each divergent pathway produces multi-stable incompatibility and is non-redundant, but the joint incompatibility of all such pathways become redundant since multiple pathways serve the same function. In the main text, we model two pathways with identical functions, each pathway contains two bi-allelic loci (first pathway: A/a & B/b, second pathway: C/c & D/d), which disrupt the pathway’s function whenever one locus becomes heterozygous. The condition for incompatibility is

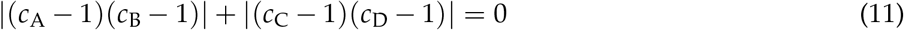

## 3. Results

### 3.1 The collapse of incompatibilities in a peripheral species

Peripheral species are small, semi-isolated populations distributed near the boundary of a large, more static species. Reproductive isolation might evolve in these lineages via elevated genetic drift and temporarily suppressed gene flow (Levin, 1970).

Consider a peripheral lineage of species 1 experiencing unidirectional gene flow from species 2. We solved the average time to collapse of incompatibilities defined by Models I & II using Equation (6) (Figure 2). The average time to collapse 〈*t_c_*〉 is defined as the first time that every individual in the peripheral lineage becomes compatible with the external species. Since hybridization is weak, population size fluctuates near *N*, the equilibrium value in the peripheral lineage. For simplicity, let *m*\* = *Nm* and *r*\* = *Nr* be the population rates of migration and recombination following classical population genetic notions.

**Figure 2:**
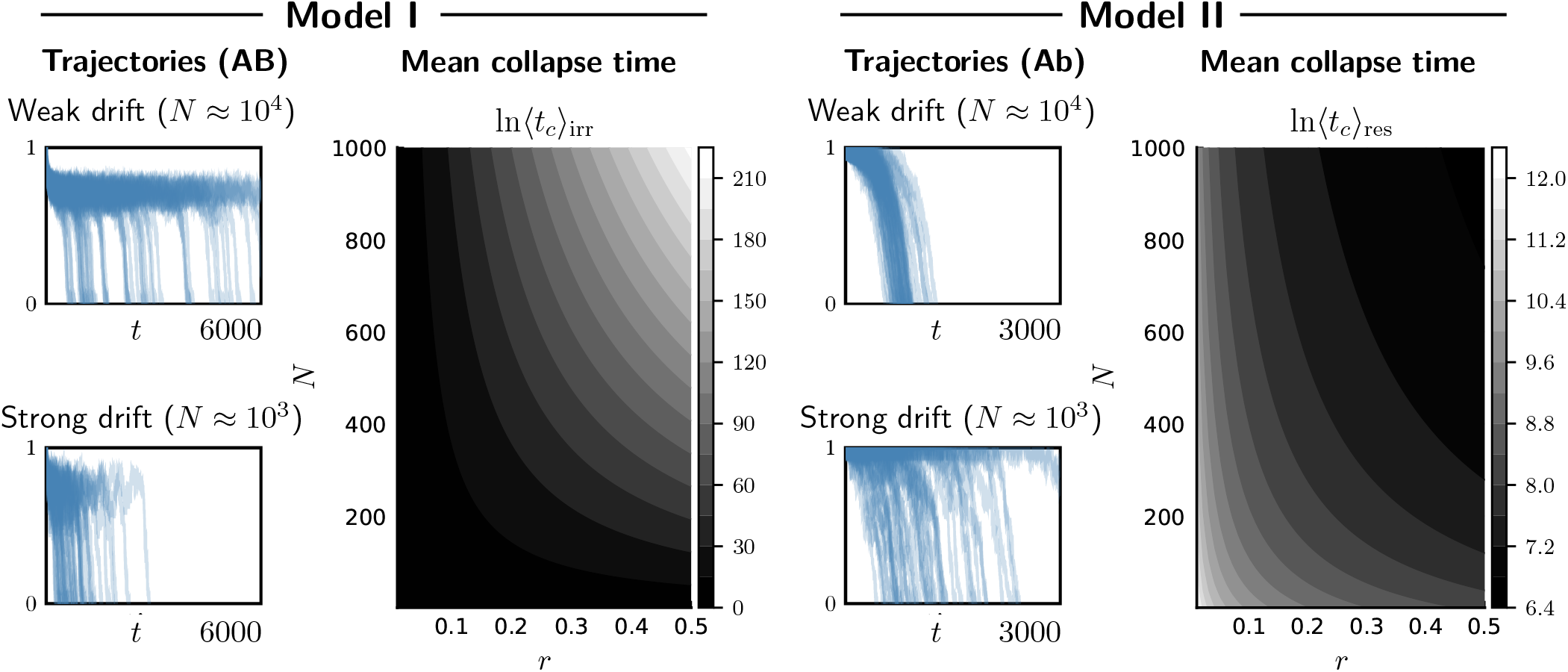
The collapse in a peripheral lineage. All trajectories are results of 50 repeated simulations of the birth-death process. Simulations have the following recombination rates: *r* = 0.055 for Model I, and *r* = 0.25 for Model II. Numerically calculated mean collapse times 〈*t_c_*〉_res_ and 〈*t_c_*〉_irr_ using Equations (13) and (12) are shown as heatmaps. All data in this figure correspond to fixed parameters: *s* = 10, *α* = 1, *m* = 0.01.

For the non-redundant model (I), its average time to collapse is approximately (see Materials and Methods: Analysis of Model I in a peripheral lineage)

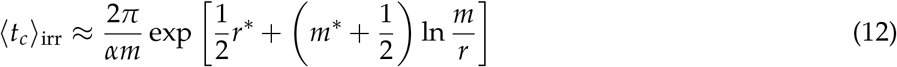

〈*t_c_*〉_irr_ approaches infinity when noise magnitude goes to zero (i.e., *N* → ∞). Here, the effect of noise depends on population migration rate *m*\* as well as population recombination rate *r*\*.

For the redundant model (II), its average time to collapse in the peripheral lineages is given by (see Materials and Methods: Analysis of Model II in a peripheral lineage)

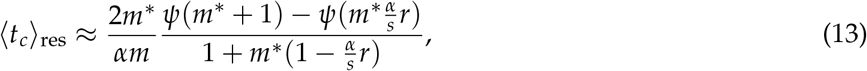

where *ψ*(·) represents the digamma function. 〈*t_c_*〉_res_ is bounded as a function of noise magnitude. As discussed before, the effect of noise is mediated only through population migration rate *m*\*.

Although a peripheral lineage represents only the simplest population structure, it is sufficient to illustrate the main difference of collapse between the two classes of incompatibilities. While redundant incompatibilities collapse continuously (bounded 〈*t_c_*〉), non-redundant ones collapse abruptly in a way similar to the shifting-balance process (Wright et al., 1932) (exponential 〈*t_c_*〉). Such dichotomy will extend further when we consider their collapse in spatially-extended populations.

There are other contrasting features in both models. In Model I, first-generation hybrids are normal, while backcross generations are largely unfit. In Model II, fitness reduction occurs immediately in first-generation hybrids, but dissipate in backcross generations. Consequently, noise affects the persistence of incompatibilities in opposite ways. In Model I, incompatibility alleles are deterministically stable in the peripheral lineage when hybridization is weak. Therefore, weaker noise will strengthen the performance of the deterministic barrier. In Model II, however, stronger noise helps create a first-generation barrier to collapse, which works by increasing the likelihood of losing the entire first-generation hybrids before they reproduce. When noise is weak, it is increasingly likely to have some first-generation hybrids surviving the incompatibility and generating backcrosses, which lead to incompatibility collapse. These effects produce almost opposite patterns of 〈*t_c_*〉 as a function of *N* (heatmaps in Figure 2).

### 3.2 The collapse of non-redundant incompatibilities in spatial systems

#### Non-redundant incompatibilities collapse via stochastic travelling waves between broadly-sympatric species

To understand the sympatric collapse of non-redundant incompatibilities, consider a coarse-grained spatial system, where two layers of demes represent local populations of two species (Figure 3A). Each deme is in one of the two discrete states: +1 and −1, representing the two parental genotype distributions **g**_1_ and **g**_2_. With nearest-neighbor migration within layers (*m*′), and hybridization between layers (*m*), each deme flips its state with a fixed rate depending on the states of its immediate neighbors. A contact probability can thus be defined for each deme as the fraction of individuals being exchanged with demes of the opposite states, which captures the net influence of other genotypes. For each deme, its rate of flipping increases with a larger contact probability. Such simplified model is a reasonable approximation to the full system, as long as the full system is bi-stable and the collapse occurs abruptly with a fixed rate. We also define a stochastic travelling wave in the discrete system as a biased movement of the boundary between two states. Since such movement requires stochastic flipping of discrete states, wave fronts propagate stochastically similar to a biased random walk. Due to the requirement of bias, the spatial boundary between opposite states needs to be asymmetric for travelling waves to exist.

**Figure 3:**
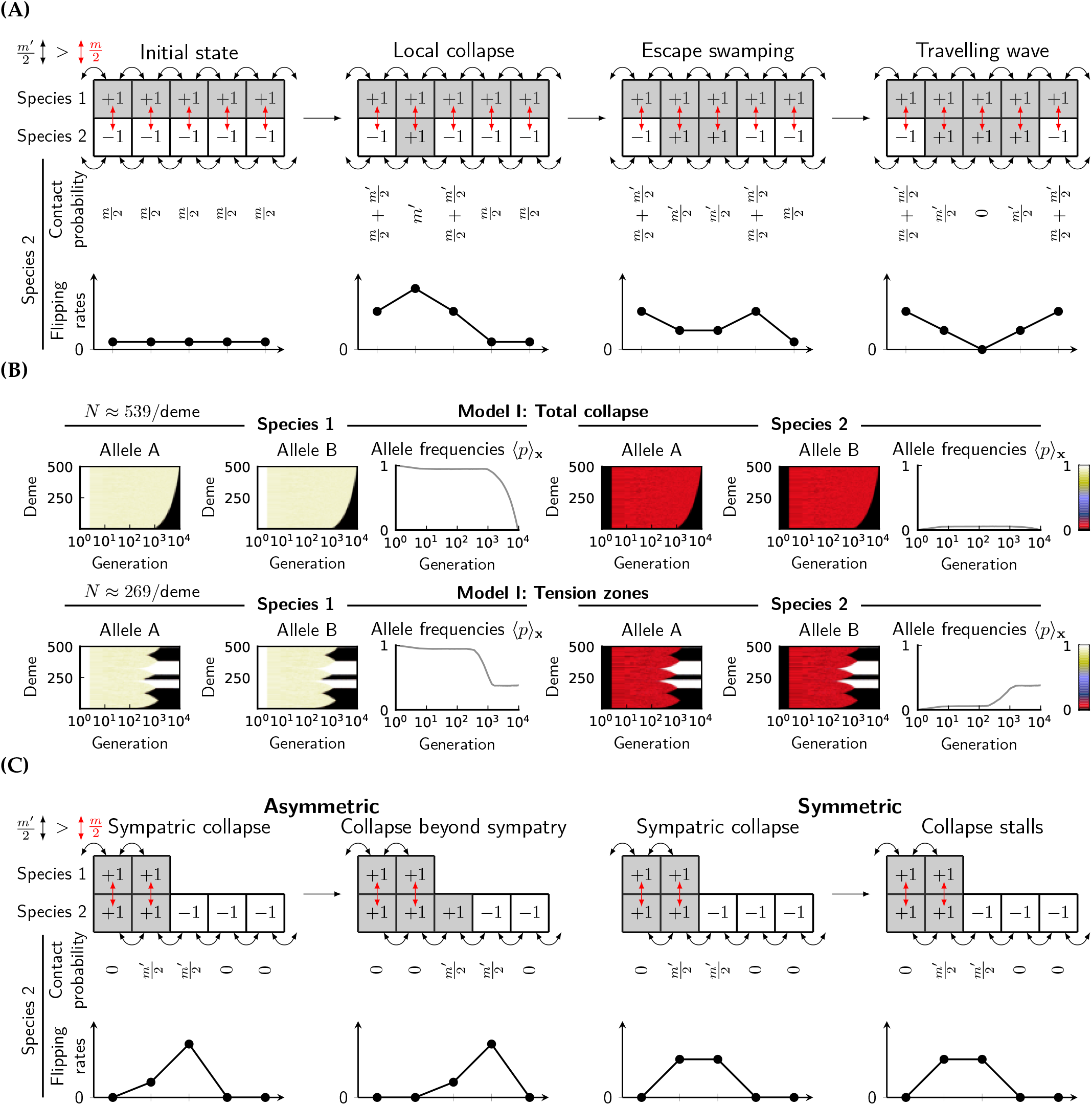
Non-redundant incompatiblities elicit travelling waves in spatial systems. **(A)** The coarse-grained system of Model II between sympatric species. Once the initial state is broken by a local collapse, the collapsed deme has to survive swamping. If it escapes swamping, the rate of flipping is higher towards the exterior of the collapsed region, which drives the propagation of travelling waves. **(B)** Two examples of SLiM simulations on Model II showing that symmetric incompatibilities could collapse completely or form multiple tension zones. Allele frequencies were taken every five generations (i.e., Δ*t* = 5 in Equation 14). Hybridization probability *m* = 0.05, and nearest-neighbor migration probability *m*′ = 0.1. **(C)** Coarse-grained systems beyond sympatry. If the two states are asymmetric, collapse extends to demes away from the hybrid zone. If the two states are symmetric, collapse stalls when it reaches the border of hybrid zones, and travelling waves cannot propagate beyond sympatry.

In general, the first event to occur is a local collapse (assimilation of discrete states) between a pair of sympatric demes. Following this event, there are two sources of asymmetry in the system which could contribute to the formation of travelling waves. First, if non-redundant incompatibilities are intrinsically asymmetric (i.e., states +1 and –1 are not exchangeable without altering the dynamics), travelling waves can form due to migration within each species (Barton and Turelli, 2011; Geldhauser and Kuehn, 2020). Second, continuing hybridization creates asymmetry that favors the wave to propagate in the same direction as the initial collapse. The second source of asymmetry means that even a completely symmetric bi-stable incompatibility can produce travelling waves, which is impossible in the absence of hybridization.

Analyzing the coarse-grained system of Model I is sufficient to show how travelling waves originate. In this symmetric model, there are three steps to initiate a travelling wave (Figure 3A). The system first waits for an initial local collapse, after which the collapsed deme has a fixed probability of not being swamped by gene flow from its neighboring con-specific demes. Finally, boundaries between the two states in the same species will move stochastically, with a bias towards expanding the collapsed region, and this is the stage where stochastic travelling waves can be observed. We simulated Models I in a one-dimensional habitat with bidirectional hybridization using software SLiM (see Materials and Methods: Simulation in SLiM). Note that SLiM implements discrete populations and discrete generations. Simulated results (Figure 3B) demonstrate the existence of travelling waves, and the final state of the entire system varies from a total collapse to the formation of multiple tension zones. The likelihood of these outcomes clearly depends on the number of initiated waves, as well as the direction of collapse at each wave front, both of which depend on the details of incompatibilities and the level of noise in each deme. The presence of travelling waves alters the time-scale of incompatibility’s persistence. On the one hand, if the initial collapse is slow, but travelling waves form readily, a total collapse between the two species is likely to occur. On the other hand, if initial collapse occurs frequently in space, and the incompatibility is symmetric, long-term tension zones will form, which segregate space into regions fixed for different genotypes.

#### Only asymmetric non-redundant incompatibilities collapse via stochastic travelling waves outside hybrid zones

When two species hybridize in sympatry, non-redundant incompatibilities can collapse via stochastic travelling waves regardless of their symmetry, but only asymmetric incompatibilities produce travelling waves beyond hybrid zones. The lack of travelling waves in symmetric systems is the result of the absence of hybridization outside hybrid zones, which leaves only the intrinsic asymmetry of incompatibilities as the driving force of travelling waves (Figure 3C). Wave propagation mechanisms under such systems are well-established results (Barton and Turelli, 2011; Geldhauser and Kuehn, 2020). Due to reduced asymmetry, even if travelling waves exist, their wave speeds will also be reduced compared to those in sympatry.

### 3.3 The collapse of redundant incompatibilities in spatial systems

#### Redundant incompatibilities collapse into quasi-neutral spatial polymorphism between broadly-sympatric species

At a local spatial scale, the collapse of redundant incompatibilities can be viewed as two parental genotype distributions converging to a common point along some quasi-neutral path in the state space G. However, the final position of this point in the state space is not unique, because noise in the system alters the trajectories throughout the collapse. Consequently, independent realizations of the same collapse will likely have variable outcomes.

In spatially-extended populations, if habitat is large, genetic drift decouples between very distant populations (isolation by distance) (Barton et al., 2013; Wright, 1943). As a result, the immediate outcome of collapse will also differ between distant populations. Between adjacent populations, however, gene flow tightly couples their genotype distributions, and no adjacent populations should be incompatible, otherwise they will resolve into compatible states. Two consequences can thus be predicted:

- A transect of genotype distributions across space, following the collapse, is also part of a quasi-neutral path in the state space.
- No local tension zones.

A signature of quasi-neutrality is that allele-frequency changes will be dominated by genetic drift. Quantitatively, across a small time interval (Δ*t*), the mean-squared allele-frequency-change across space in each species 〈|Δ*p*|^2^)_x_ will follow:

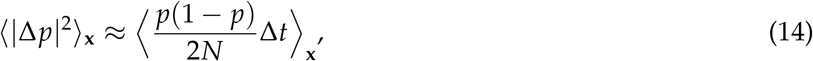

where 〈·〉_**x**_ represents averaging across all local populations of that species.

To demonstrate quasi-neutrality, we simulated redundant Models II, III & IV in a one-dimensional habitat with bidirectional hybridization across 500 demes in SLiM. As shown in Figure 4, all systems have a phase of rapid collapse following the onset of hybridization, after which sympatric local populations share similar allele frequencies between different species. However, in the long-term, allele frequencies still vary substantially across space, and the overall frequencies in the entire species change very slowly. When comparing the mean-squared response (〈|Δ*p*|^2^〉_x_) against neutral genetic drift (Δ*t* = 5 generations), we found that the log-ratio of quantities on the two sides of Equation (14) deviates from zero in the short-term, but fluctuates around zero in the long-term, confirming that genetic drift has dominated allele frequency changes after the initial collapse. This marks the existence of a long quasi-neutral phase. As all alleles involved in incompatibilities are purged at very low rates in the quasi-neutral phase, it is possible for con-specific samples from different local populations to be incompatible. Further, different inbred lines generated from a single local populations could also yield different levels of incompatibility when crossed with another species, because local polymorphism is tolerated, and is not subject to strong selection.

**Figure 4:**
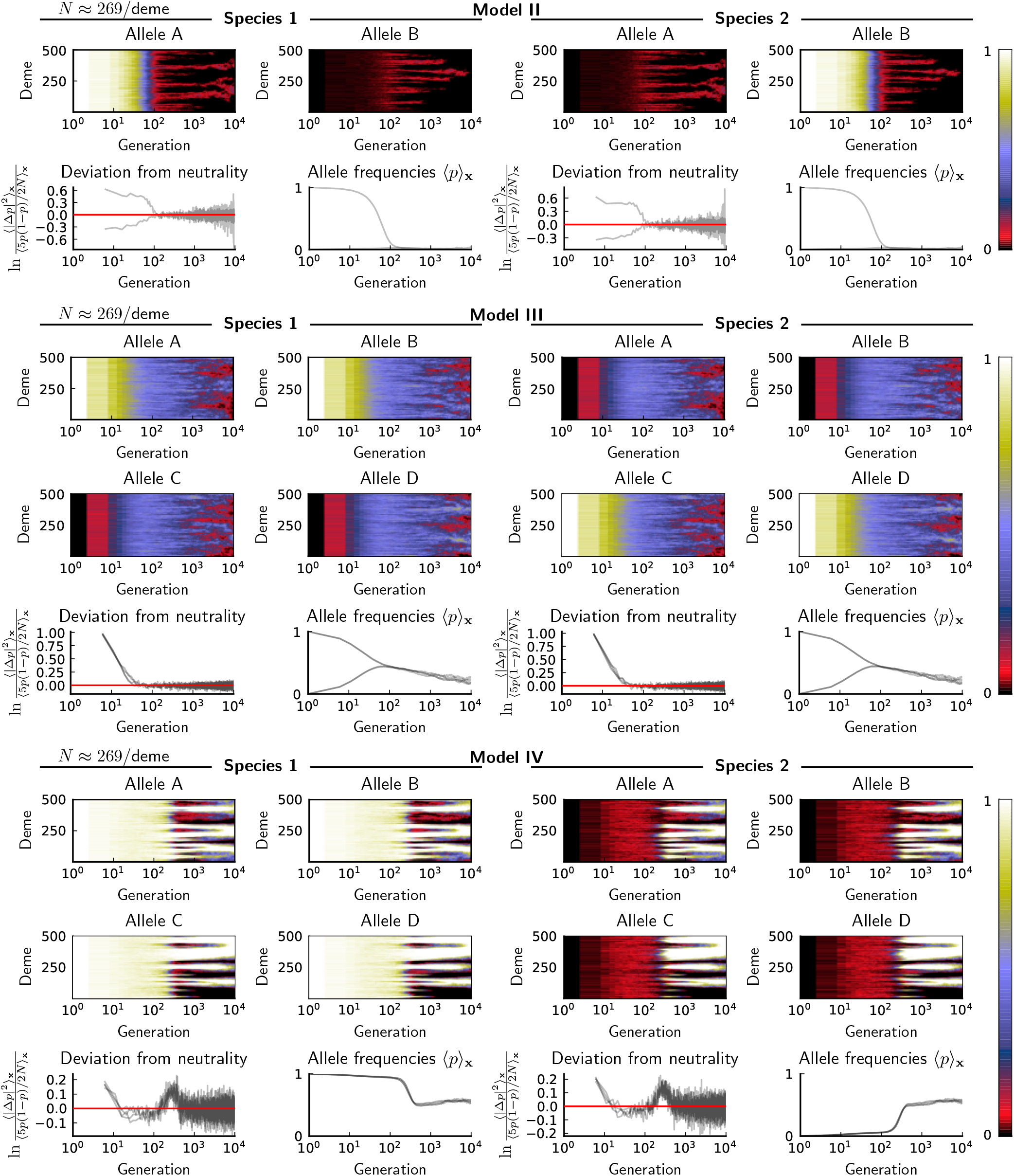
SLiM simulations demonstrating that redundant incompatibilities collapse into quasi-neutral polymorphism between broadly-sympatric lineages. Heatmaps are allele frequencies across space and time. Allele frequencies were taken every five generations (i.e., Δ*t* = 5 in Equation 14). For all three models, hybridization probability *m* = 0.05, and nearest-neighbor migration probability *m*′ = 0.1. In deviation from neutrality, red lines are expectations under neutral genetic drift.

Spatial allele frequencies in the quasi-neutral phase closely follow the topology of quasi-neutral paths in the state space. In the redundant multi-stable system (Model IV), despite a lack of tension zones, alleles from different pathways still segregate into large spatial blobs, and fix in an alternating order in space. In the redundant uni-stable system (Model III), the pseudogenized alleles are seldom fixed locally, and their average frequencies decreases through time; However, their functional homologs are fixed in an alternating order in space similar to the redundant multi-stable system. The fact that functional homologs or functional pathways are fixed in an alternating order in space is congruent with the aforementioned prediction relating a spatial transect of genotype distributions to a quasi-neutral path: in both systems, quasi-neutrality requires at least one functional component to be fixed locally.

#### The collapse of redundant incompatibilities is limited by spatial dispersal outside hybrid zones

We have already shown that redundant incompatibilities collapse into global quasi-neutrality between broadly-sympatric lineages. A similar quasi-neutrality argument can be applied beyond hybrid zones because incompatibility alleles are purged indirectly there: the collapse outside the hybrid zone is driven only by the dispersal of compatible genotypes, which gradually replace parental genotypes throughout space.

Consequently, the collapse rate of incompatibility alleles outside a hybrid zone is simply bounded above by the accumulation rate of compatible genotypes via spatial dispersal. Formally, suppose an incompatibility allele is initially unique to one species and is fixed. At time *t* post hybridization, let 〈*p*〉 (**x**, *t*) be its expected frequency in that species at location **x** in the geographic range D. Let *p_s_*(**x**, *t*) be the probability that a randomly sampled lineage, at location **x** ∈ D and at time *t*, has never hit the hybrid zone throughout its genealogical history. Then we have

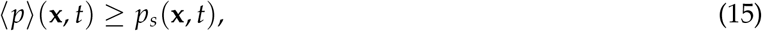

In general, *p_s_*(**x**, *t*) is the survival probability of a spatial dispersal process, with immediate killing near the boundary of hybrid zones. Although the exact form of *p_s_*(**x**, *t*) depends on dispersal mechanisms and habitat geometry, we assume a simple model to demonstrate Equation (15). Let D be interval [0, *L*] in one species, and suppose hybridization only affects the left boundary *x* = 0. Further, assume population density is uniform and dispersal is driven by the standard diffusion with variance *σ*^2^ per unit time (*σ*^2^ = *m*′ in discrete populations with nearest-neighbor migration). Then:

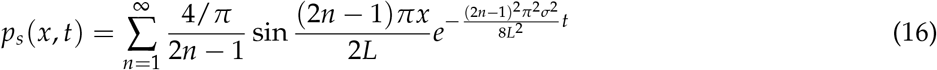

Using this formula, at the right-most boundary *x* = *L*, the time it takes on average to decrease allele frequencies to *p* is lower-bounded by

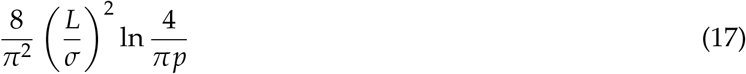

The term *L*^2^/*σ*^2^ signifies a general pattern in one-dimensional dispersal by standard diffusion, that the rate of collapse away from the hybrid zone slows down at least quadratically with the habitat size, which is a consequence from the scaling relationship in standard diffusion (displacement 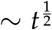). By this logic, dispersal characteristics will affect the collapse rate away from the hybrid zone profoundly. If long-range dispersal occurs (i.e., a heavy-tail dispersal kernel), the collapse will be much quicker than that under standard diffusion, because incompatibility alleles are brought into the hybrid zone more quickly to be selected out. Conversely, lineages with restricted dispersal (can be modelled with subdiffusion) will collapse much more slowly away from the hybrid zone.

The bound *p_s_* is compared with simulated results for Models II, III & IV in a one-dimensional habitat with a very narrow hybrid zone (Figure 5). Despite the complexity of simulated systems, *p_s_* curves robustly predict the “envelope” of allele frequency clines, and the ensemble-average 〈*p*〉 of all allele frequency clines are always bounded by *p_s_*.

**Figure 5:**
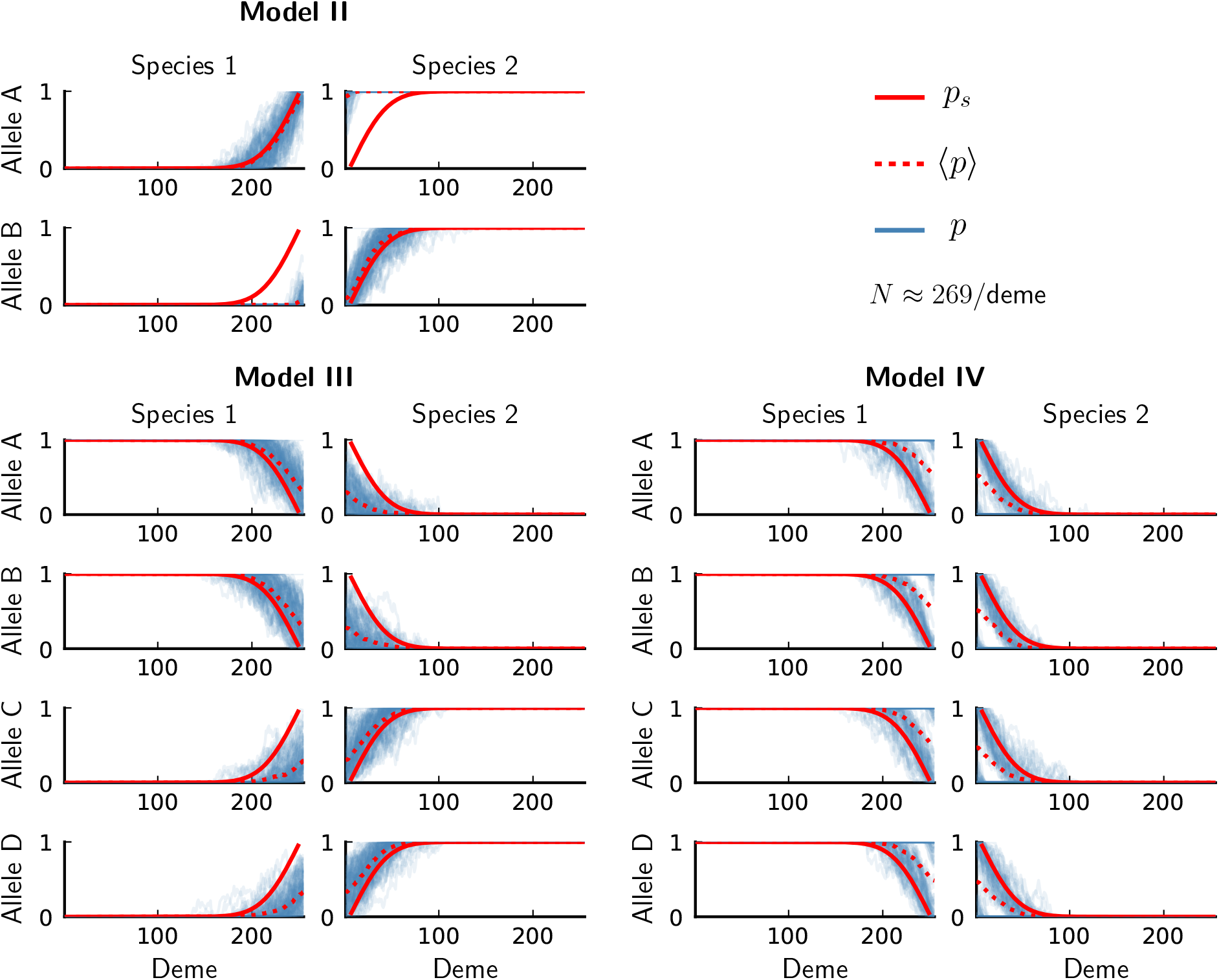
Simulations in SLiM of Models II, III & IV. Each model was run for 100 realizations, and allele frequencies (*p*, blue curves) were taken at generation *t* = 9901. The hybrid zone has five consecutive demes, and there are 250 demes forming a one-dimensional array outside the hybrid zone in each species. Hybridization probability *m* = 0.05, and nearest-neighbor migration probability *m*′ = 0.1. Solid red lines are the survival probability *p_s_* calculated using Equation (15) with *σ*^2^ = *m*′. Dashed red lines are the ensemble-average 〈*p*〉 of allele frequencies *p*.

## 4. Discussion

Hybridization enables incompatibility genes to interact in natural populations. Although multilocus epistasis appears formidably complex to derive general conclusions of interacting incompatibilities in nature, we demonstrate that useful information can be extracted from the level of redundancy in incompatibility mechanisms. The key to the generality of classification based on genetic redundancy is that many empirical incompatibility systems seem to have evolved via duplications of genes or pathways, followed by developmental system drift in each isolated lineages.

In summary, we have shown that different levels of genetic redundancy could lead to qualitatively different behaviors of incompatibilities under hybridization. When hybridization is a weak force, redundant incompatibilities change continuously in time, often associated with various quasi-neutral behaviors. However, non-redundant incompatibilities can be understood using bi-stable (or multi-stable) dynamics, which produce abrupt changes in allele frequencies (shifting-balance, travelling waves, tension zones, etc.), and will be more sensitive to noise such as genetic drift. In existing literature, redundant incompatibilities, such as classical Dobzhansky-Muller systems, are considered unable to sustain hybridization unless maintained by ecological selection (Bank et al., 2012). From our point of view, while redundancy renders the system unstable to gene flow, it also serves as a buffer against complete extinction of incompatibility alleles, and the likelihood of wide-spread polymorphism could be high with redundant incompatibilities in spatially-extended populations.

This framework ignores external (ecological) selection, thus the dynamics are solely determined by the intrinsic compatibility between divergent genomes. In principle, it is straightforward to augment the analysis with external selection, which will likely break the symmetry among genotypes in our models and will favor particular genotypes in a particular species, thus decreasing the likelihood of collapse. Nonetheless, the types and forms of external selection are less predictable across different taxonomic groups, and environmental change will frequently alter the landscape of external selection such that species maintained by external selection could still collapse (Taylor et al., 2006; Vonlanthen et al., 2012). However, this ecological route to species collapse represents a completely different regime outside the scope of this paper.

We should emphasize that hybridization is not the only route to incompatibility variation. For instance, different mutations arising in different local populations could contribute to variable incompatibilities across space. Nonetheless, the classification based on genetic redundancy will also be useful in analyzing patterns not produced via hybridization, because a higher redundancy will always lead to greater tolerance of polymorphic incompatibility alleles. As in the spatial simulations discussed before, such variation could also follow quasi-neutral dynamics, which evolves slowly through time.

## 5. Materials and Methods

### 5.1 Construction of dynamics from a birth-death process

For a focal diploid monoecious population, define the following continuous-time birth-death process with three types of events:

- Generate a single offspring from a random bi-parental mating within the focal population (birth)
- Generate a single offspring from a hybrid mating with a different species (birth)
- Eliminate an existing individual (death)

The rate of each process is described as follows. Let *n_j_* be the number of individuals of genotype *j*, so that **n** = (*n*_0_, *n*_1_,…) is a state under the birth-death process. Each monoecious individual independently reproduces with rate *α*. If an individual reproduces, it randomly selects a partner from the focal population with probability 1 – *m*, or from the foreign with probability *m*. After selecting the partner, a single offspring from the bi-parental mating is added to the focal population. Apart from reproduction, each existing individual of genotype *j* dies independently with rate *μ*_0_ + *β*∑*_j_n_j_* + *s_j_*, where *μ*_0_ is the normal mortality rate, *β* controls density-dependent mortality, and *s_j_* controls viability selection. Let *R_j_*(**n**) and *M_j_*(**n**) be the overall birth rate and death rate of genotype *j* in the focal population:

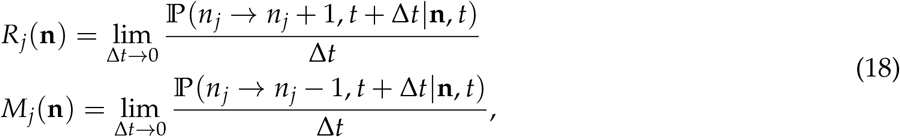

Let *Q*(*j*′ × *j*″ → *j*) be the probability of generating an offspring of genotype *j* from a mating between parents of genotypes *j*′ and *j*″, then

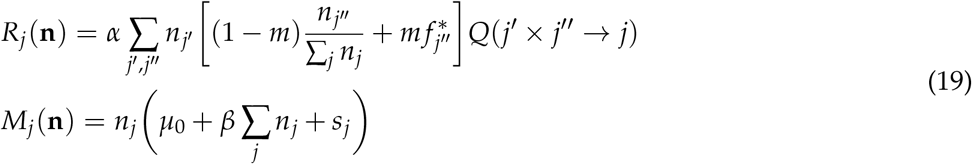

Hence, the entire process is defined as the following continuous-time 1-step Markov jump process

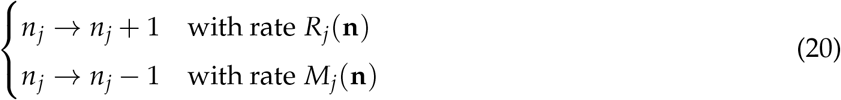

The transition density *p_t_*(**n**) = *p*(**n**, *t*|**n**′, *t*′) of the birth-death process can be approximated around **n** to the second order by the following Fokker-Planck equation:

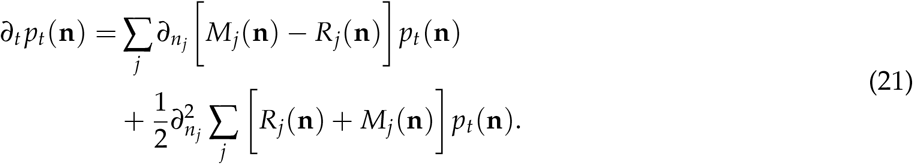

Let **Q** be the stoichiometric matrix with *Q*(*j*′ × *j*″ → *j*) at entry (*j*′ × *j*″, *j*). Let ***μ*** be the mortality rate vector with entries *μ*_0_ + *β*∑*_j_ n_j_* + *s_j_*. Let **g** = **n**/∑*_j_ n_j_* be genotype frequencies. The convection part of Fokker-Planck equation can be transformed to the following differential equations between **g**_1_ (focal species) and **g**_2_ (foreign species):

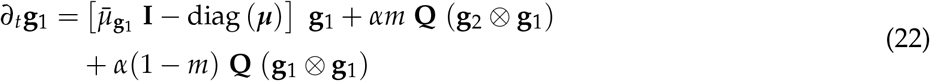

The diffusion part of Fokker-Planck equation will be used in approximating the mean collapse time in Models I & II.

### 5.2 Separation of time-scales

Without loss of generality, we show the procedure under the scenario of redundant incompatibilities. It can be modified for non-redundant incompatibilities by grouping **f**_2_ with **f**_3_ and following the same steps. In the main text, Equation (22) is written succinctly as:

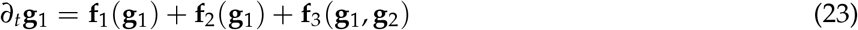

Since **f**_3_(**g**_1_, **g**_2_) is relatively slow, during Δ*t* time it perturbs the system from **g** to **g**\* = **g** + **f**_3_Δ*t*, which is within the vicinity of the equilibrium set 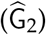 of equation *∂*_*t*_**g**_1_ = **f**_1_(**g**_1_) + **f**_2_(**g**_1_). Such proximity allows for the linearization of dynamics:

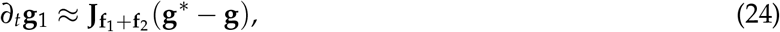

where **J**_**f**_1_+**f**_2__ is the Jacobian of **f**_1_ + **f**_2_ near **g**. This linear dynamics occur on a fast time-scale, so the joint perturbation from both slow and fast parts is

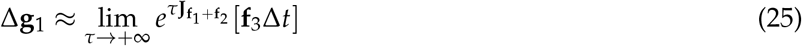

Taking Δ*t* → 0 yields the final equation (5). Note that such perturbative approach is accurate only when the fast part brings the state sufficiently close to the equilibrium set 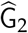 (deviation 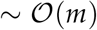).

### 5.3 Analysis of Model I in a peripheral lineage

For Model I, the direction of gene flow is from species 2 to species 1 (aabb → AABB), and we track the frequency of haplotype AB (denoted as *θ*_AB_) since recombined haplotypes are at very low frequencies. Since the time-scale of collapse is at least inversely-scaled with the rate of gene flow, we could re-scale time by the rate of gene flow (i.e., *τ* = *tαm*/2) to simplify calculation. By transforming to the frequency of haplotype AB and working out the asymptotic equation by separation of time-scales defined above, we have:

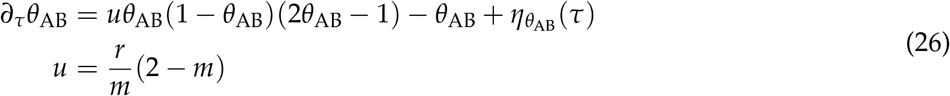

where *η*_*θ*_AB__ (*τ*) is a white noise following

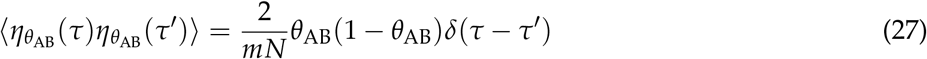

This dimensional reduction technique produces reasonably accurate dynamics of haplotype frequencies in both Models I and II (Figure 7).

It is straighforward to show that the stochastic process evolves along a potential function proportional to *U*(*θ*_AB_) = *uθ_AB_*(1 – *θ*_AB_) – ln(1 – *θ*_AB_). The potential function defines a metastable state at *θ*_AB_ = *θ*_*_, and a saddle state at *θ*_AB_ = *θ*_**_. *θ*_AB_ quickly approaches *θ*_*_ at the onset of hybridization, so we may directly calculate the mean escape time from *θ*_AB_ = *θ*_*_ to *θ*_AB_ = 0 as the mean collapse time of the incompatibility. Define *Ψ*(*θ*_AB_) = exp[– *mNU*(*θ*_AB_)], the mean escape time is then

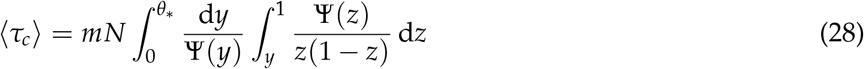

If the effect of genetic drift is weak (*mN* ≫ 1), the outer integrand Ψ(*y*)^−1^ is sharply peaked at *θ*_**_, and the inner integrand 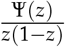 is peaked to the right of *θ*_**_ at *θ*_*_. Hence the inner integral is approximately constant when *y* is being integrated near *θ*_**_. We may re-write the expression as

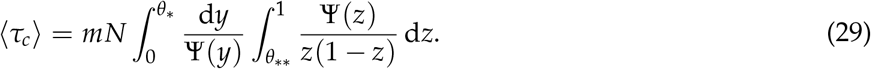

Let

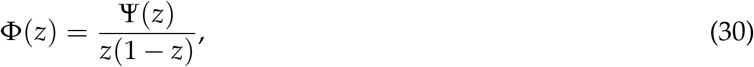

and

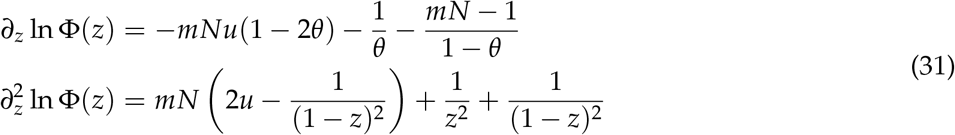

*θ*_*_ is thus the first solution smaller than 1 to the equation *∂_z_* ln Φ = 0. We use a Beta approximation to both Φ and Ψ^-1^:

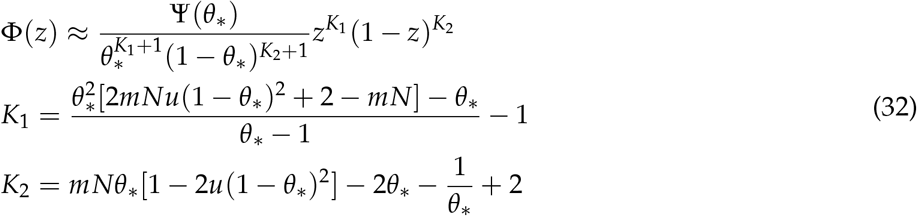

Thus

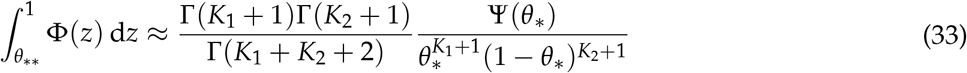

The function Ψ^-1^ has the following expansion:

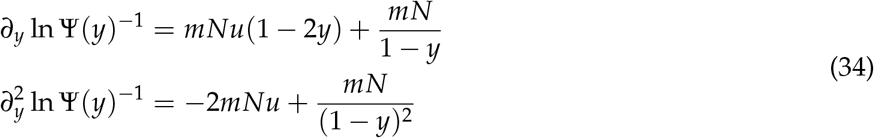

From the first equation we know 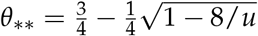. Then

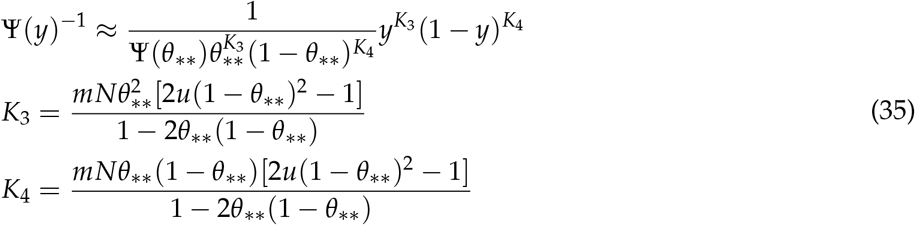

Thus

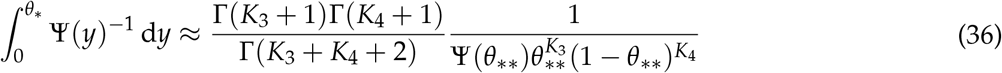

Combining both, we have

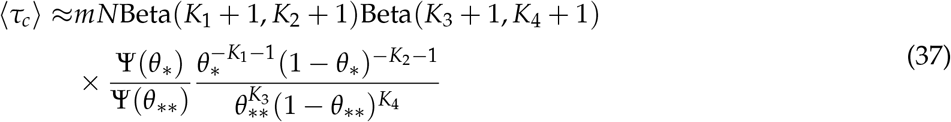

For even larger *mN*, we may substitute the Beta approximation by a Gaussian. The Gaussian approximation to Φ(*z*) near *θ*_*_ is

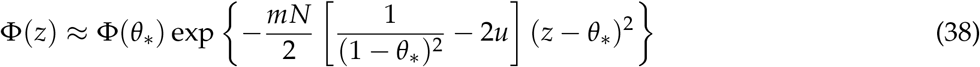

Thus,

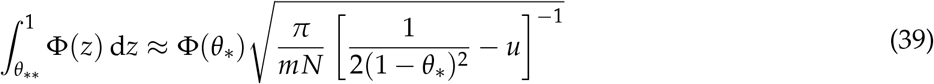

Similarly,

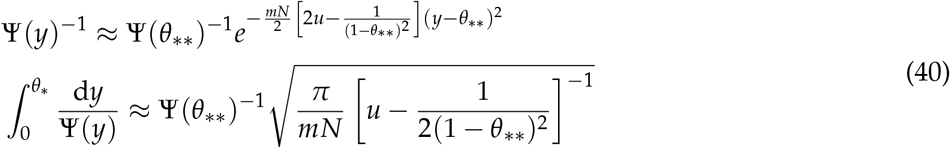

Finally,

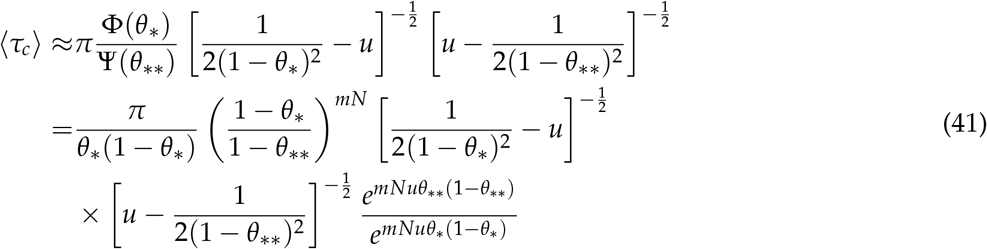

*θ*_*_ and *θ*_**_ are approximately the two non-zero fix points of the deterministic part of Equation (26) in the large *mN* limit:

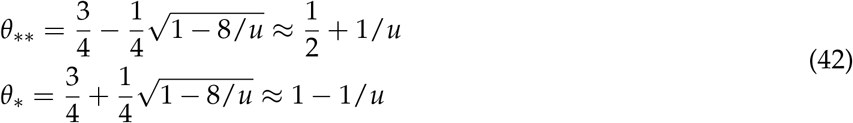

Substituting into the expression gives the following final approximation:

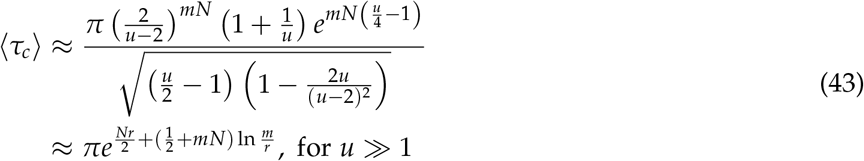

converting to the original time by 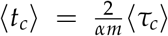 yields Equation (12). Both approximations of 〈*t_c_*〉 were checked against simulated birth-death processes in Figure 6B, and we found that simulated 〈*t_c_*〉 lies between the two approximations.

**Figure 6:**
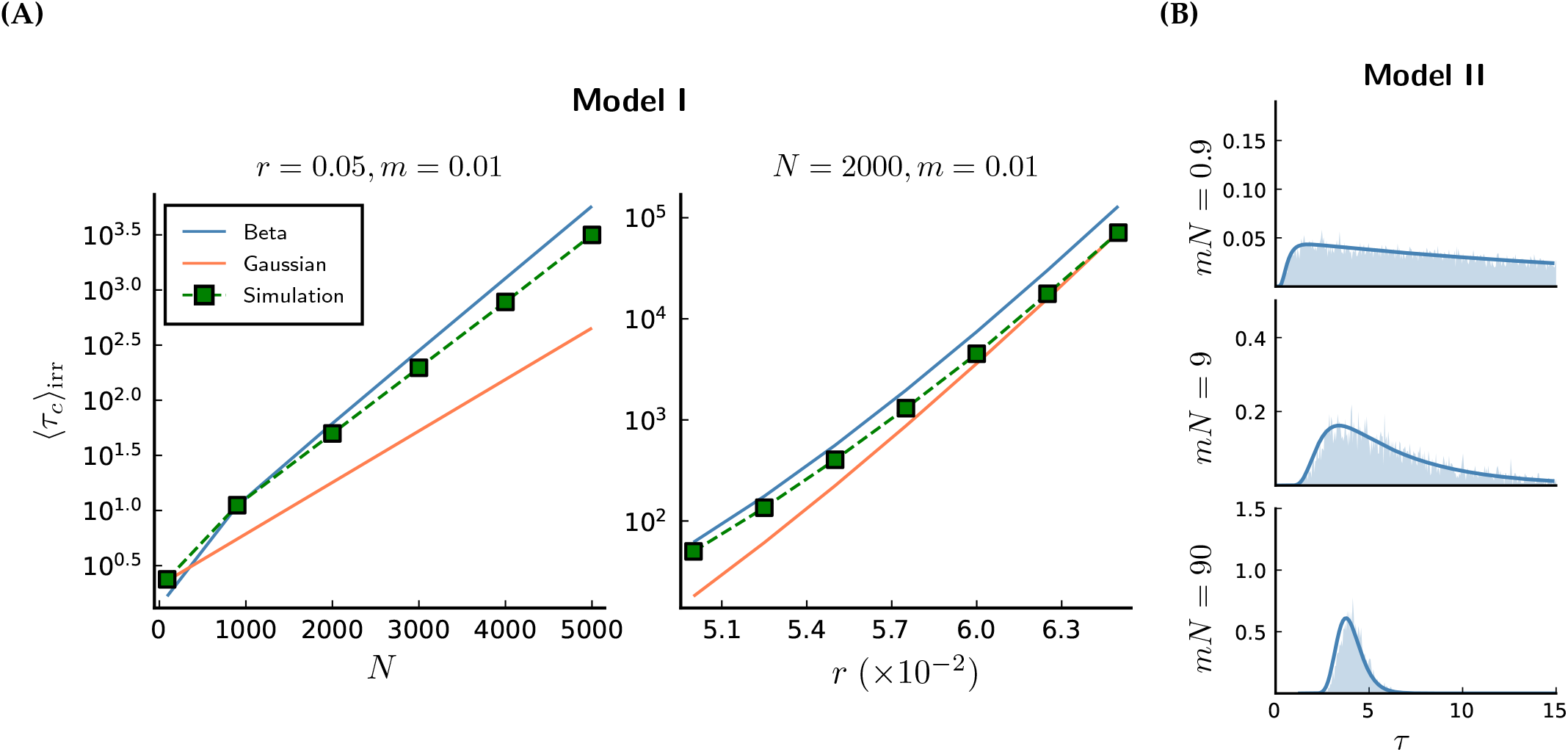
Comparison of the collapse time in Models I & II between analytic approximations and simulated birthdeath prcesses. For each combination of parameters we simulated 3000 replicates of the entire birth-death process. The equilibrium population size *N* is allowed to vary by adjusting the density-dependent mortality rate parameter *β*. These parameters are fixed: *α* = 1, *μ* = 0.1, *s* = 10, and *m* = 0.01. **(A)** The approximate mean collapse time 〈*t_c_*〉_irr_ of Model I versus simulated results. **(B)** The approximate distribution of *t_c_* in Model II using Equation (45) (solid curves) versus simulated results (histograms). Recombination probability *r* = 0.25.

**Figure 7:**
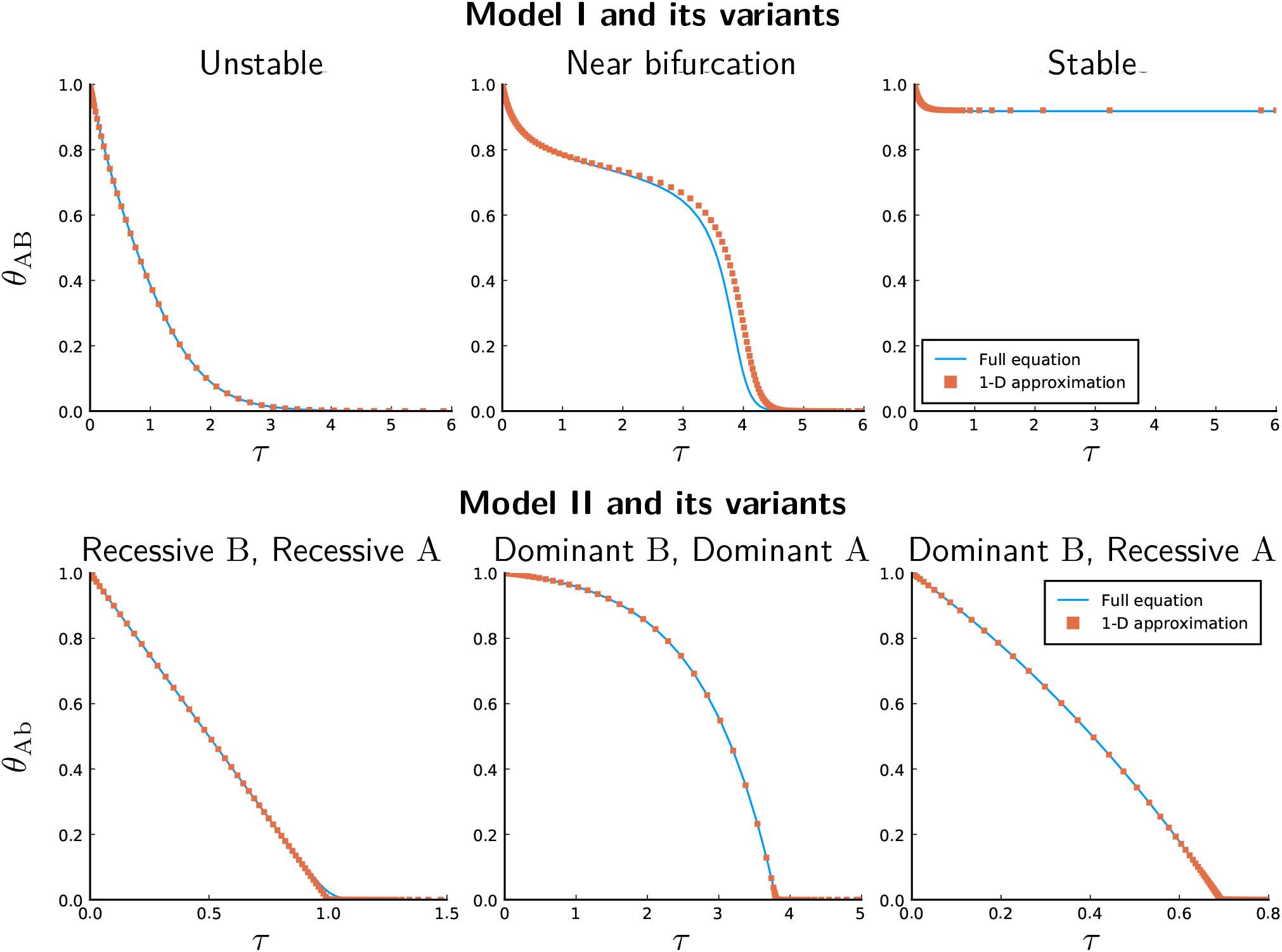
One-dimensional asymptotic equations like Equations (26) and (44) are good approximations to the full haplotype dynamics under strong incompatibilities and weak hybridization in Models I & II and their variants. The figure shows the deterministic solutions to the high-dimensional ODEs of haplotype dynamics under the birth-death model in a peripheral lineage (blue curves), together with the one-dimensional approximations using time-scale separation techniques (orange squares). Model I parameters: *s* = 10, *α* = 1, *μ*_0_ = 0.1, *m* = 10^−2^, *r* = 0.01 (Unstable), *r* = 0.075 (Near bifurcation), *r* = 0.15 (Stable). Model II parameters: *s* = 10, *α* = 1, *μ*_0_ = 0.1, *m* = 10^−5^, *r* = 0.25.

### 5.4 Analysis of Model II in a peripheral lineage

For Model II, the direction of gene flow is from species 2 to species 1 (aaBB → AAbb). Haplotype AB in species 1 is always negligibly small since it always causes incompatibilities, and so we could track the frequency of haplotype Ab (denoted as *θ*_Ab_) to determine the collapse dynamics. The asymptotic equation of *θ*_Ab_ in the re-scaled time *τ* = *tam*/2 is

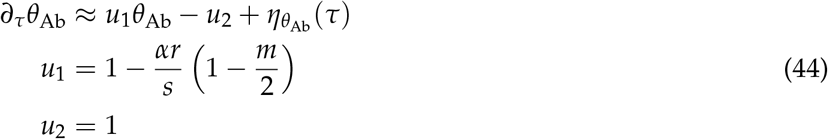

The noise satisfies Equation (27).

Let the collapse time *τ_c_* be the first time that *θ*_Ab_ = 0. Using eigenfunction expansion, we obtained the distribution of *τ_c_*:

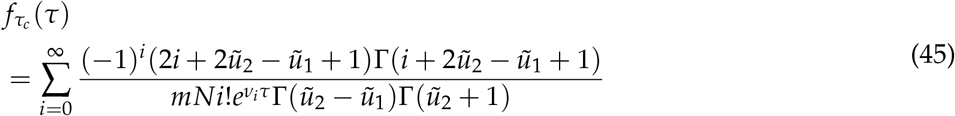

where

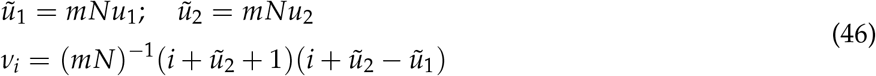

We can check the high accuracy of our approximation by simulating the original birth-death process of Model II and compare the true distribution of *τ_c_* versus Equation 45 (Figure 6B).

The mean collapse time 〈*τ_c_*〉 satisfies

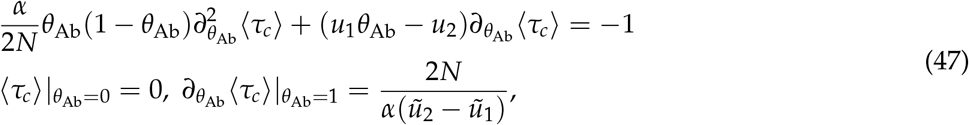

which has the general solution

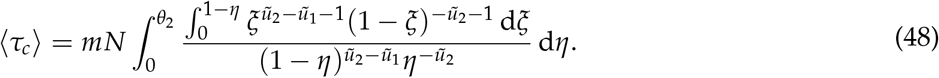

Solving the integral gives the following specific solution, where *ψ* is the digamma function and _3_*F*_2_ is the generalized hypergeometric function:

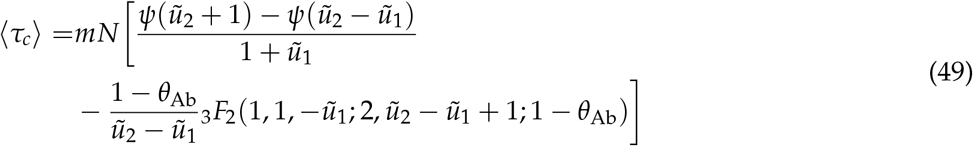

At *θ*_Ab_ = 1, the second term becomes 0 and we have the mean collapse time of a population initial fixed with AAbb:

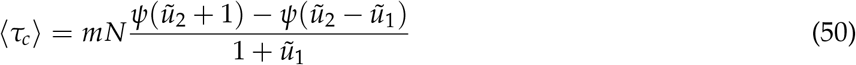

Transforming back to the original time-scale using 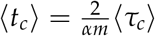 yields the result in Equation (13).

### 5.5 Simulation methods

#### Simulating the birth-death process

The full birth-death process was simulated in Julia-1.5 using the Gillespie algorithm implemented in the package `DifferentialEquations.jl`. For a large population, the process is approximated by the tau-leaping algorithm, where during a fixed small time interval [*t*, *t* + Δ*t*], the numbers of jumps of each type are drawn from Poisson distributions using the rates at time *t*, and the process leaps forward to *t* + Δ*t* by updating the variables using the numbers of jumps.

#### Simulation in SLiM

SLiM-3.6 (Haller and Messer, 2019) was used to simulate spatial models. Since it uses discrete generations and discrete demes, it only approximates the birth-death process. However, important qualitative results still hold. We implemented the following non-Wright-Fisher generation cycles in SLiM:

- Each population of size N randomly draws a Poisson number of offsprings of mean 1.5*N* to produce.
- For each offspring, randomly draw parent 1 from the current population. For parent 2, draw with probability *mN*_foreign_/(*mN*_foreign_ + (1 – *m*) *N*) from the foreign species, otherwise draw from the current population. Cross parent 1 with parent 2 and add the offspring to the offspring pool.
- After offspring production, remove all parents, and each offspring dies with probability specified by the form of viability selection.
- Migrate survived offsprings within each species following designated population structures. Each offspring independently migrates with probability *m*′. If it migrates, it randomly chooses a nearest-neighbor deme as the destination.

For viability selection, the probability of survival for each offspring is the product between the probability of surviving density-dependent selection (exp(–*N*/*K*)) and the probability of surviving selection-by-incompatibility (Model I: compatible = 1, incompatible = 0; Models II, III, IV: compatible = 1, incompatible = 0.25). *K* controls the population size.

## 6. Data availability

The code for simulation is available at https://github.com/tzxiong/2020_The_collapse_of_genetic_incompatibilities

## 7. Acknowledgments

T.X. is partially funded by the NSF-Simons Center for Mathematical and Statistical Analysis of Biology at Harvard (award number #1764269) and the Harvard Quantitative Biology Initiative during the project. We thank Prof. John Wakeley for his detailed comments on several drafts of the manuscript. We thank Prof. Robin Hopkins and Prof. Naomi Pierce for providing useful general feedbacks. We thank Nathaniel Edelman and Neil Rosser for the discussion on *Heliconius* incompatiblities that inspired the current work. We also thank Fernando Seixas, Miriam Miyagi, Yuttapong Thawornwattana, Sarah Dendy and three anonymous reviewers of a previous version of the manuscript.

## Footnotes

1 To one extreme, if the whole system is completely redundant, its dynamics will be equivalent to neutral evolution, where any path in the state space can be Γ_2_

